# Innervation density governs crosstalk of GPCR-based norepinephrine and dopamine sensors

**DOI:** 10.1101/2024.11.23.624963

**Authors:** Ricardo C. López, Natalie Noble, Özge D. Özçete, Xintong Cai, Gillian E. Handy, Jonathan W. Andersen, Tommaso Patriarchi, Yulong Li, Pascal S. Kaeser

## Abstract

GPCR-based fluorescent sensors are widely used to correlate neuromodulatory signaling with brain function. While experiments in transfected cells often reveal selectivity for individual neurotransmitters, sensor specificity in the brain frequently remains uncertain. Pursuing experiments in brain slices and in vivo, we find that norepinephrine and dopamine cross-activate the respective sensors. Non-specific activation occurred when innervation of the cross-reacting transmitter was high, and silencing of specific innervation was indispensable for interpreting sensor fluorescence.

## Main

G protein-coupled receptor (GPCR)-based fluorescent sensors have revolutionized our means of studying neuromodulatory signaling in the brain. Their ability to discriminate neurotransmitters relies on the properties of the respective endogenous GPCRs that are used for sensor engineering. Studies describing new sensors typically measure EC50 values in transfected cells by comparing responses to locally puffed neurotransmitters at increasing concentrations^1–6^. While this approach generally finds high specificity, it is often unclear how this in vitro selectivity translates to the brain. In addition to sensor affinities, local innervation densities of neuromodulatory axons and the organization of release and reuptake machinery may strongly influence which transmitters are reported by a given indicator. This is particularly relevant for transmitters with similar chemical structures.

We examined the selectivity of GPCR-based norepinephrine and dopamine sensors^2,4,5^ that are widely used, often in brain areas innervated by both transmitter systems (Supplemental text). We first quantified the innervation densities of these transmitters in primary motor cortex (M1) and dorsal striatum. Prior work has found dense dopamine innervation in dorsal striatum and prominent presence of norepinephrine axons in M1 cortex^7–11^. To measure axonal densities, we genetically labeled dopamine axons by Cre-dependent expression of synaptophysin-tdTomato in DAT^IRES-Cre^ mice (Fig. 1a-c, Supplemental fig. 1). We prepared brain slices containing dorsal striatum and M1 cortex and performed antibody staining against RFP and the norepinephrine transporter (NET) to visualize dopamine and norepinephrine axons, respectively (Fig. 1b+c).

**Figure 1.**
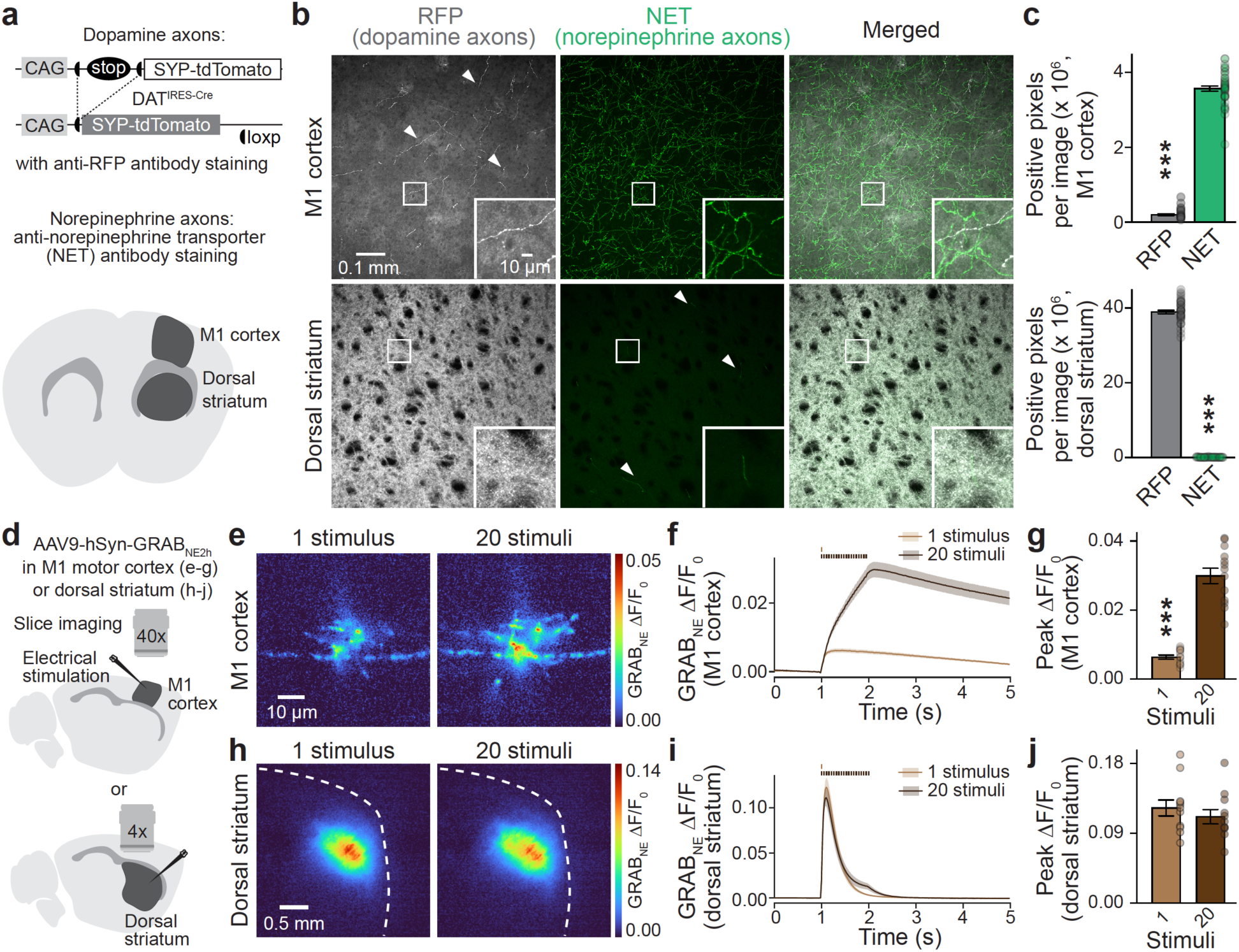
Cortex and striatum have distinct dopamine and norepinephrine innervation and GRAB_NE_ dynamics. **a.** Strategy for quantification of dopamine and norepinephrine innervation with confocal microscopy. Dopamine axons were labeled by anti-red fluorescent protein (RFP) antibody staining in coronal brain slices of DAT^IRES-Cre^ mice with Cre-dependent synaptophysin-tdTomato (SYP-tdTomato) expression. Norepinephrine axons were labeled with anti-norepinephrine transporter (NET) antibodies. **b, c.** Representative images of M1 cortex and dorsal striatum (b) and quantification of RFP and NET staining (c) as the total number of positive pixels in image stacks after binarization. In b, brightness and contrast in the gray (RFP) channel were differentially adjusted to render cortical dopamine axons visible and to prevent saturation of dorsal striatum; M1 cortex 40 slices from 8 mice, dorsal striatum 40/8. **d.** Schematic of imaging in acute sagittal slices containing M1 cortex and dorsal striatum. Wide- field fluorescence imaging was performed 3 weeks after stereotaxic AAV injections of the corresponding brain areas to express GRAB_NE_. Electrical stimulation was used to elicit fluorescence transients. **e-j.** Representative images of peak GRAB_NE_ fluorescence in response to 1 stimulus or 20 stimuli at 20 Hz in M1 cortex (e) or dorsal striatum (h) and quantification of GRAB_NE_ ΔF/F0 time series (f, i) and peak ΔF/F0 (g, j); M1 cortex 13 slices from 3 mice, dorsal striatum 11/3. Data are mean ± SEM; *** p < 0.001, as assessed by: two-tailed Mann-Whitney rank-sum test in c, two-tailed Wilcoxon signed-rank test in g. For assessment of somatic stainings in LC and SNc, see Supplemental fig. 1. For pharmacological blockade of GRAB_NE_ transients in brain slices, see Supplemental fig. 2. For assessment of GRAB_NE_ in HEK293T cells, see Supplemental fig. 3.

Confocal images revealed segregation of the two fluorescence signals within each area. In M1 cortex, norepinephrine fibers predominated, and dopamine fibers were sparse. Conversely, dopamine axons were dense in dorsal striatum while norepinephrine axons were infrequent. Quantification of each signal suggested that M1 cortex contains ∼15 times more norepinephrine than dopamine innervation, and dorsal striatum as much as ∼350 times more dopamine than NET-stained innervation (Fig. 1c). Assessment of staining in the locus coeruleus (LC) and substantia nigra pars compacta, the corresponding sites of norepinephrine and dopamine neuron somata, confirmed the specificity of the genetic strategy (Supplemental fig. 1).

To examine the transmitter dynamics resulting from these innervation patterns, we first expressed the norepinephrine sensor GRAB_NE_2h (α2 adrenergic receptor-based, abbreviated as GRAB_NE_)^4^ in M1 cortex or dorsal striatum and analyzed evoked fluorescence transients in acute brain slices with widefield microscopy (Fig. 1d-j). Single electrical stimuli and stimulus trains (20 stimuli at 20 Hz) yielded fluorescence increases in M1 cortex consistent with norepinephrine innervation (Fig. 1e-g). These transients were fully blocked by inhibition of action potential firing but persisted in a cocktail of blockers of synaptic transmission (Supplemental fig. 2). Single stimuli induced small GRAB_NE_ transients, and 20-stimulus trains produced a buildup with a peak amplitude ∼5 times greater than that of a single stimulus (Fig. 1f+g), similar to prior electrochemical studies^9,12,13^. Robust GRAB_NE_ transients were also detected in the dorsal striatum despite the near absent norepinephrine innervation, though with different properties (Fig. 1h-j). Peak striatal GRAB_NE_ transients were similar between single and train stimuli, and they decayed more rapidly than the transients in M1 cortex.

The distinct innervation patterns and GRAB_NE_ signals in M1 cortex and dorsal striatum (Fig. 1) suggest that GRAB_NE_ might report dopamine in the striatum. This is further supported by the similarity of striatal GRAB_NE_ transients (Fig. 1h-j) to the properties of dopamine release^14–16^. The initial characterization of GRAB_NE_ found an EC50 of ∼190 nM for norepinephrine and ∼9 μM for dopamine^4^. Correspondingly, the GRAB_NE_ response in transfected cells was ∼10-fold greater for 10 μM norepinephrine puffs compared to 10 μM dopamine puffs (Supplemental fig. 3).

Dopamine-laden vesicles, though, have an intravesicular dopamine concentration of up to ∼1 M, and the dopamine concentration at the pore of a fusing vesicle^9,17,18^ far exceeds commonly used concentrations for probing selectivity^1–6^.

To directly test whether dopamine induces GRAB_NE_ transients in dorsal striatum, we used mice with conditional deletion of the release site-organizer protein RIM in dopamine neurons (RIM cKO^DA^, Fig. 2a+b). In these mutants, evoked dopamine release is disrupted when assessed with amperometry, whole-cell electrophysiology, or GRABDA . Striatal slices from RIM cKO^DA^ mice displayed strong reductions in evoked GRAB_NE_ transients compared to RIM control slices in response to both single and train stimuli (Fig. 2c-e). The same was true when nicotinic acetylcholine receptors, which trigger dopamine release via induction of dopamine axonal action potentials^21–23^, were blocked (Supplemental fig. 4). We next used nLightG, an alternate GPCR-based norepinephrine sensor engineered from the α1 adrenergic receptor^5^. Like GRAB_NE_, nLightG also reported striatal dopamine release (Fig. 2f-h), though with several fold lower overall amplitudes. RIM cKO^DA^ slices had ∼90% reduced nLightG responses, a reduction similar to that observed with GRAB_NE_.

**Figure 2.**
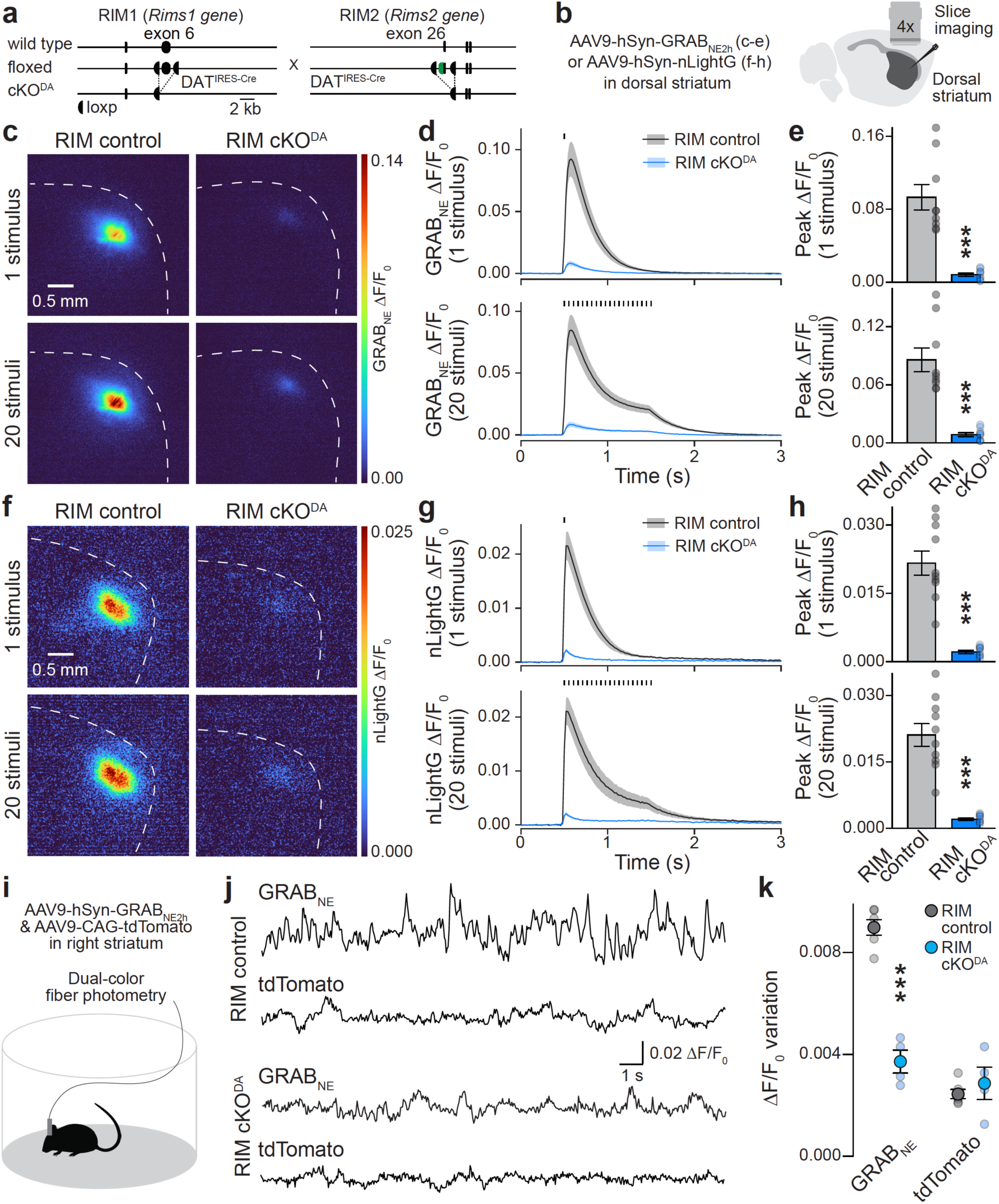
Norepinephrine sensors detect dopamine in dorsal striatum. **a.** Strategy for conditional deletion of RIM1 and RIM2 in dopamine neurons as established before^14,16,19,20^. **b.** Schematic of slice imaging in dorsal striatum after expressing GRAB_NE_ or nLightG. **c-e.** Representative images of peak GRAB_NE_ fluorescence in response to 1 stimulus or 20 stimuli at 20 Hz (c) and quantification of GRAB_NE_ ΔF/F0 time series (d) and peak ΔF/F0 (e); RIM control 9 slices from 3 mice, RIM cKO^DA^ 9/3. **f-h.** As in c-e, but with nLightG instead of GRAB_NE_; RIM control 10/3, RIM cKO^DA^ 10/3. **i.** Schematic of fiber photometry in freely moving mice co-expressing GRAB_NE_ and tdTomato in the right dorsal striatum. **j, k.** Representative examples of GRAB_NE_ and tdTomato ΔF/F0 (j) and fluorescence variation quantified as standard deviation (SD) of GRAB_NE_ and tdTomato ΔF/F0 (k); RIM control 6 mice, RIM cKO^DA^ 4. Data are mean ± SEM; *** p < 0.001, as assessed by: two-tailed Mann-Whitney rank-sum tests in e and h, two-tailed unpaired t-test in k. For assessment of striatal GRAB_NE_ transients while blocking nicotinic acetylcholine receptors, see Supplemental fig. 4. For post-hoc assessment of fiberoptic cannula positions, see Supplemental fig. 5.

To test whether the cross-reactivity is also present in vivo and detected in response to endogenous neural activity, we performed fiber photometric GRAB_NE_ recordings in freely moving RIM control and RIM cKO^DA^ mice (Fig. 2i-k, Supplemental fig. 5). Fluorescence transients were measured as mice explored an open field arena, and we quantified GRAB_NE_ fluctuations as variation of ΔF/F0. GRAB_NE_ fluctuations were readily detected in RIM control mice, but nearly absent in RIM cKO^DA^ mice (Fig. 2j+k). This is similar to striatal transients measured with dopamine sensors^16^. In summary, these data establish that two commonly used norepinephrine sensors are almost exclusively activated by dopamine in the dorsal striatum.

Given the robust cortical norepinephrine innervation, we next examined the ability of GRAB_NE_ and nLightG to detect norepinephrine in M1 cortex. We first established that chemical lesion of norepinephrine neurons in the right LC with 6-hydroxydopamine (6-OHDA)^24^ abolished cortical NET staining in the same hemisphere (Fig. 3a-c, Supplemental Fig. 6). We then bilaterally expressed GRAB_NE_ or nLightG in M1 cortex in mice with unilateral LC lesions (Fig. 3d+h). Both sensors showed increased fluorescence in response to single and train stimuli and in each case, the transients were strongly decreased when the LC was lesioned (Fig. 3d-k). Importantly, striatal GRAB_NE_ responses were not impaired after LC lesion (Supplemental fig. 6c-f), indicating that they did not arise from LC neurons and that dopamine neurons were not lesioned by 6- OHDA injection into the LC.

**Figure 3.**
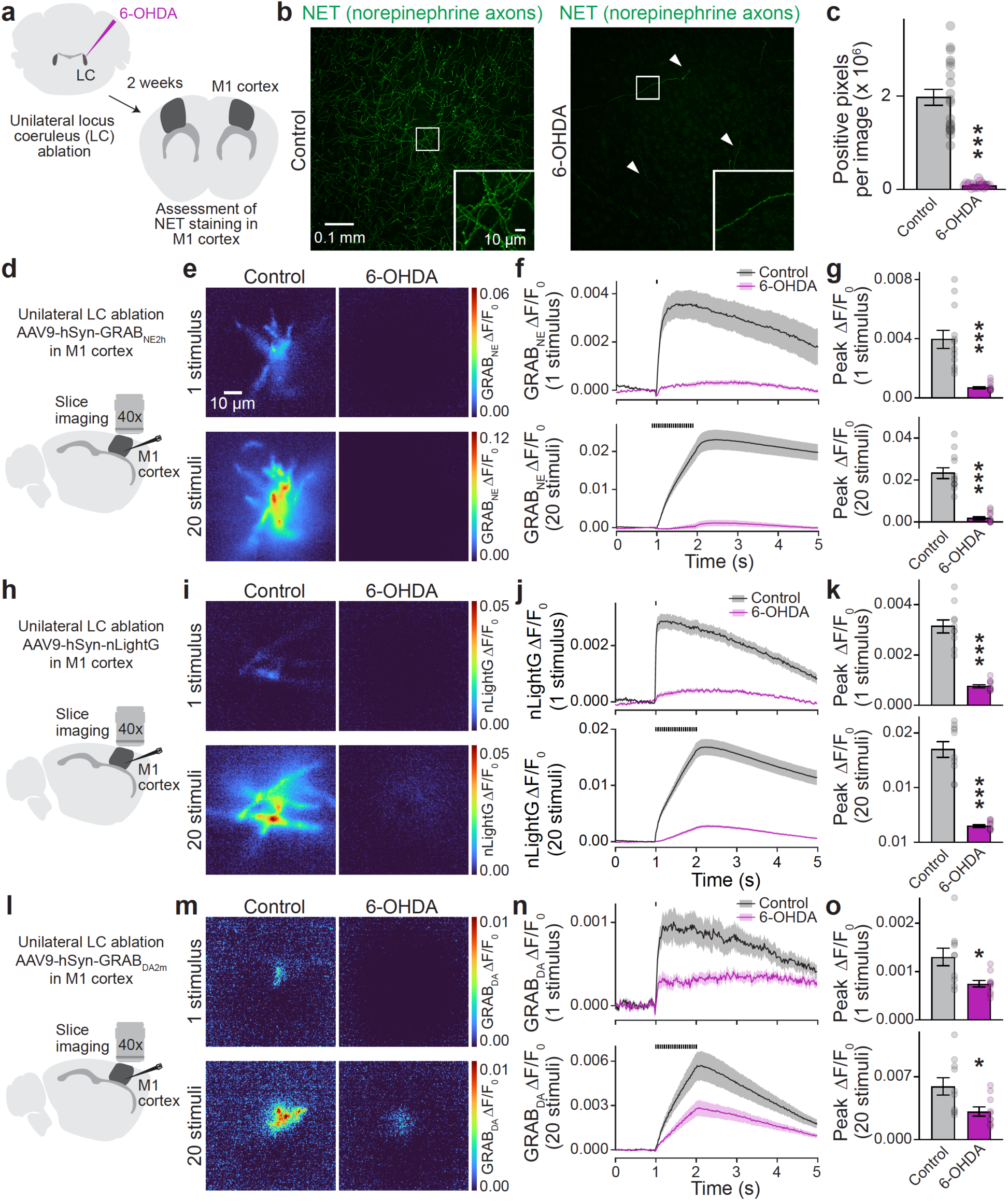
Norepinephrine and dopamine sensors in M1 cortex. **a.** Schematic of unilateral LC ablation with 6-OHDA. Two weeks after injection, norepinephrine axons were labeled with anti-NET antibodies. Analyses were performed on confocal images of coronal sections in contralateral (control) and ipsilateral (6-OHDA) hemispheres. **b, c.** Representative images of M1 cortex (b) and quantification of NET signals (c); 20 slices from 4 mice. **d.** Schematic of bilateral slice imaging in M1 cortex expressing GRAB_NE_ following unilateral ablation of the right LC. **e-g.** Representative images of peak GRAB_NE_ fluorescence in response to 1 stimulus or 20 stimuli at 20 Hz in ipsilateral (6-OHDA) and contralateral (control) hemispheres (e) and quantification of GRAB_NE_ ΔF/F0 time series (f) and peak ΔF/F0 (g); control 12 slices from 3 mice, 6-OHDA 12/3. **h-k.** As in d-g, but with nLightG instead of GRAB_NE_; control 10/3, 6-OHDA 10/3. **l-o.** As in d-g, but with GRABDA instead of GRAB_NE_; control 10/3, 6-OHDA 10/3. Data are mean ± SEM; *** p < 0.001, * < 0.05 as assessed by: two-tailed Mann-Whitney rank-sum tests in c, g, k, and o (1 stimulus); two-tailed unpaired t-test in o (20 stimuli). For assessment of histology and GRAB_NE_ responses in striatum following 6-OHDA injection in LC, see Supplemental fig. 6. For assessment of GRABDA in HEK293T cells, see Supplemental fig. 7. For assessment of GRAB_NE_ and GRABDA transients in LC, see Supplemental fig. 8.

Given the structural similarities of dopamine and norepinephrine, do dopamine sensors detect norepinephrine? We tested GRABDA2m, a commonly used D2 receptor-based sensor (abbreviated as GRABDA) with an EC50 of ∼80 nM and ∼1.2 μM for dopamine and norepinephrine, respectively^2^. In transfected HEK293T cells, puffing 10 µM dopamine or norepinephrine produced similar GRABDA fluorescence changes (Supplemental fig. 7). We have previously shown that striatal GRABDA transients are strongly impaired in RIM cKO^DA^ mice^16^, establishing that GRABDA reports dopamine in the striatum. When we monitored GRABDA signals in M1 cortex (Fig. 3l-o), transients induced by single and train stimuli were impaired by ∼50% following 6-OHDA lesion of LC. Hence, GRABDA responds at least partially to norepinephrine in M1 cortex. The remaining transients may be due to cortical dopamine innervation, which may be enhanced upon LC ablation^24^, or to other transmitters.

The LC receives hardly any dopamine innervation (Supplemental fig. 1)^10^. To evaluate transmitter dynamics in the LC, we expressed either GRAB_NE_ or GRABDA bilaterally in the LC and monitored fluorescence transients. Evoked LC GRAB_NE_ transients had similar characteristics (Supplemental fig. 8a-d) to those measured in cortex (Fig. 1d-g), and so did LC GRABDA transients (Supplemental fig. 8e-g). With both sensors, single pulse transients were small compared to those produced by 20 stimuli at 20 Hz, and the transients had prolonged decays. These findings suggest that GRABDA and GRAB_NE_ report norepinephrine in LC.

The presented results establish that GPCR-based norepinephrine and dopamine sensors have prominent crosstalk in the brain. Although in vitro affinity measurements reflect sensor properties, a key determinant of sensor activation in the brain is the ratio of axonal innervation densities of specific modulators, which can vary by 100-fold or more between different brain areas (Fig. 1). Our findings are important for many past and ongoing studies (Supplemental text). Typically, sensor transients are interpreted as reporting the transmitter they were named after, and appropriate loss-of-function experiments are often not performed. A commonly used control to verify sensor activation is pharmacological sensor inhibition, though this approach does not validate transmitter specificity. We propose that crosstalk should always be experimentally addressed in studies using GPCR-based sensors for neuromodulatory transmitters. To this end, genetic or chemical silencing are essential to interpret sensor signals in brain slices and in vivo. These manipulations should be done under the specific conditions used in a study and are a necessary standard in the future.

The cross-reactivity we observe for dopamine and norepinephrine sensors is unlikely limited to these two transmitters. Many modulators, including serotonin, other monoamines, and more than a hundred neuropeptides operate in the brain^9^, and these systems might cross-activate a variety of sensors. Achieving full signal selectivity is particularly challenging based on the structural similarities of some of these transmitters.

Synthesis pathways might further complicate the interpretation of catecholamine sensor data. For example, neurons synthesize norepinephrine through enzymatic conversion of dopamine, and LC neurons might co-release dopamine and norepinephrine^25,26^. Thus, the sensors might report mixed signals in response to activities of norepinephrine neurons when enzymatic conversion is incomplete.

Finally, we note that cross-talk is not a “flaw” of the sensors but instead reflects the properties of GPCRs. Dopamine and adrenergic receptors, for example, can bind both dopamine and norepinephrine^27–29^. Receptors for other neuromodulators also interact with several ligands^30–32^. While this non-selectivity complicates interpretations of sensor imaging and fiber photometry, it underscores that receptor crosstalk likely represents a biologically meaningful mechanism to convey information and to regulate cells, circuits, and behavior.

## Acknowledgements

We thank Fabiola Urena Ortiz and Vanessa Charles for technical support, and J. Williams, J. Lebowitz and members of the Kaeser laboratory for discussions and/or comments on the manuscript. This work was supported by grants from the NIH (R01NS103484, R01DA056109, R01NS083898 and R01DA058777 to P.S.K.), a Harvard Medical School Neurobiology Spark Grant (to P.S.K.), a Harvard Brain Initiative Bipolar Disorder Seed Grant (to P.S.K.), and a Harvard Medical School Goldenson Research Fund Award (to P.S.K.). R.C.L. is the recipient of an NSF graduate research fellowship (DGE2140743). O.D.O. is the recipient of Human Frontier Science Program postdoctoral fellowship. J.W.A. is the recipient of a Stuart H.Q. & Victoria Quan fellowship. X.C. is currently at the Beth Israel Deaconess Medical Center. We acknowledge the Neurobiology Imaging Facility and the Core for Imaging Technology & Education at Harvard Medical School.

## Author contributions

Conceptualization: R.C.L. and P.S.K.; Methodology: R.C.L., N. N., O.D.O., X.C., J.W.A.; Resources: X.C., G.E.H., T.P., Y.L.; Investigation: R.C.L., N.N.; Formal Analysis: R.C.L., N.N.; Writing-Original Draft: R.C.L. and P.S.K.; Writing - Review and Editing: R.C.L., N.N., O.D.O., X.C., J.W.A., G.E.H., T.P., Y.L., P.S.K.; Supervision: P.S.K.; Funding Acquisition: P.S.K.

## Methods

### Mice

For morphological experiments, DAT^IRES-Cre^ mice (RRID:IMSR_JAX:006660, B6.SJL- Slc6a3^tm1.1(cre)Bkmn^/J)^33^ were bred to mice containing a CAG promoter-driven loxP-STOP-loxP- synaptophysin-tdTomato cassette (SYP-tdTomato) in the *Rosa26* locus (RRID: IMSR_JAX:012570, B6;129S-Gt(ROSA)26Sor^tm^^34.1(CAG-Syp/tdTomato)Hze^/J) as established before^14^. For conditional gene knockout experiments, mice carrying floxed alleles for RIM1 (RRID:IMSR_JAX:015832, Rims1^tm3Sud^/J)^34^ and RIM2 (RRID:IMSR_JAX:015833, Rims2^tm1.1Sud^/J)^35^ were bred to DAT^IRES-Cre^ mice^33^ as previously established^14,16,19,20^. RIM cKO^DA^ mice were homozygous for floxed RIM1 and RIM2 and heterozygous for DAT^IRES-Cre^. RIM control mice were heterozygous for both floxed RIM alleles and DAT^IRES-Cre^ and were either paired littermates of the experimental mice or age-matched non-littermate mice from intercrosses of the same alleles. Mice used for non-mutant comparisons were either wild type mice or contained floxed RIM1 and RIM2 alleles but were negative for DAT^IRES-Cre^. Male and female mice were included in all experiments regardless of sex. Intracranial injections for viral expression were performed at 5 to 12 weeks of age while functional and morphological analyses were performed at 8 to 17 weeks of age. Mice were maintained on a mixed background (containing C57BL/6 and 129/Sv) and housed in a room set to 20 to 24 °C and 50% humidity.

Mice used for fiber photometry were housed in a reversed light-dark cycle, and experiments were performed during the dark phases. Genotype comparisons were performed by an experimenter blind to genotype during data acquisition and analyses. Experiments followed approved protocols of the Harvard University Institutional Animal Care and Use Committee.

### Confocal image acquisition and processing

Morphological analyses were adapted from protocols used previously^14,16,19,20^. Mice were deeply anesthetized with 5% isoflurane and transcardially perfused with 30 mL of phosphate-buffered saline (PBS) followed by 40 mL of 4% paraformaldehyde (PFA) in PBS. Brains were post-fixed in 4% PFA in PBS for 24 hours and stored in PBS until sectioning. Mice with stereotaxic 6- OHDA injections were perfused two weeks following the injection. 50 µm thick brain slices were prepared in ice-cold PBS using a vibratome. Antigen retrieval was performed for 30 minutes at 60 °C in a solution containing (in mM) 150 NaCl, 10 Tris Base, 1 EDTA, and 0.05% Tween 20 (pH 9.0). Slices were then permeabilized in PBS containing 0.25% Triton X-100 (PBST) in three consecutive 10-minute incubations at room temperature and blocked in PBST containing 10% normal goat serum for 2 hours at room temperature. Staining with primary antibodies was performed for 24 hours at 4 °C in PBST with 10% normal goat serum. Sections were then washed three times for 10 minutes in PBST, followed by secondary antibody staining for 2 hours at room temperature in PBST with 10% normal goat serum. After another three 10-minute washes, the slices were mounted on glass slides for imaging. The primary antibodies used were mouse IgG1 anti-NET (1:1000 Synaptic Systems #260 011, RRID:AB_2636907, lab antibody code A265), guinea pig anti-RFP (1:1000, Synaptic Systems #390 004, RRID:AB_2737052, A258), and rabbit anti-TH (1:2000, Millipore #AB152, RRID:AB_390204, A66). Secondary antibodies used were Alexa Fluor 405 goat anti-rabbit (1:500, Thermo Fisher Scientific #A- 31556, RRID:AB_221605, lab antibody code S1), Alexa Fluor 488 goat anti-mouse IgG1 (1:500, Thermo Fisher Scientific #A-21121, RRID:AB_2535764, S7), and Alexa Fluor 568 goat anti-guinea pig (1:500, Thermo Fisher Scientific #A-11075, RRID:AB_2534119, S27). Confocal images were captured on an inverted spinning disk confocal microscope with a 20x, 0.8 numerical aperture air objective. Identical acquisition settings were used within a specific channel for an entire experiment. For all regions, 30 to 50 images were acquired in z from each section at a 0.9 µm step size. 20 contiguous planes were manually selected for analyses. For quantification of axonal innervation, background subtraction in each image plane was performed using the rolling ball algorithm (with a radius of 20 pixels for M1 cortex and a radius of 50 pixels for dorsal striatum), and the resulting images were binarized using thresholding modules from Python’s Scikit-image package. In M1 cortex, both channels were binarized with the Triangle thresholding method. In the dorsal striatum, the NET staining was binarized with the Triangle method and the RFP staining with the Otsu method. The number of positive pixels in each image plane was summated to quantify the total signal. To calculate the number of double-positive RFP and TH somata in the LC and SNc, 20-plane maximum projection z-stack images were created and each channel’s brightness and contrast were identically adjusted across images. Double-positive somata were manually counted. To analyze NET signals after 6-OHDA ablation of LC, brightness and contrast were identically adjusted across images. Individual z-stack planes were binarized with the Triangle method, and the number of positive pixels was summated. Representative images for figures were processed with Fiji and are composed of 20- plane maximum intensity projection z-stacks. In Fig. 1b, brightness and contrast in the green (NET) channel were adjusted identically in M1 cortex and striatum while brightness and contrast in the gray (RFP) channel were differentially adjusted to render cortical dopamine axons visible and to prevent saturation of striatal dopamine axons. For representative images in other figures, brightness and contrast of each channel were adjusted linearly and identically.

### GRAB_NE_ and GRABDA experiments in HEK293T cells

HEK293T cells were cultured as previously established^36,37^. Specifically, cells acquired from ATCC (CRL-3216, RRID: CVCL_0063, purchased mycoplasma-free, human cell line of female origin) were expanded and stored as frozen stocks until use. After thawing, the cells were grown in filtered Dulbecco’s Modified Eagle Medium (DMEM) supplemented with 10% fetal bovine serum (Atlas Biologicals F-0500-D) and 1% Penicillin-Streptomycin. HEK293T cells were split every 1 to 3 days at a ratio of 1:3 to 1:10. Cell batches were replaced after ∼20 passages by thawing a fresh vial from the expanded stock. For puffing experiments, HEK293T cells were plated on 0.1 mm-thick Matrigel-coated glass coverslips (12 mm) at ∼10-20% confluency in 24-well plates. After ∼24 hours, cells were transfected with the calcium phosphate method at ∼50- 60% confluency with 350 ng of pCMV-GRAB_NE_2h (lab plasmid code p1109, provided by Y.L.)^4^ or 200 ng of pCMV-GRABDA2m (p1110, provided by Y.L.)^2^. Between 24 to 48 hours after transfection, individual coverslips were transferred to a recording chamber perfused with a recording solution containing (in mM) 126 NaCl, 2.5 KCl, 2 CaCl2, 1.3 MgSO4, 1 NaH2PO4, 12 glucose 26.2, and NaHCO3 at 33 to 35 °C with a flow rate of 2 to 3 mL/min. The recording solution was continuously bubbled with 95% O2 and 5% CO2. Fluorescence imaging was performed using an Olympus BX51WI epifluorescence microscope. Fluorescence signals were excited with a 470 nm LED and digitized with a scientific complementary metal-oxide-semiconductor camera (sCMOS, Hamamatsu Orca-Flash4.0). Data were acquired with a 60x, 0.9 numerical aperture water-immersion objective at 512 x 512 pixels/frame and at 20 frames/s, with an exposure time of 50 ms. Dopamine hydrochloride or noradrenaline bitartrate was dissolved in MilliQ H2O to generate 100 mM stocks. These stocks were then diluted into recording solution to generate 10 µM working solutions and puffed for a 3-second epoch onto the cells using a pulled glass pipette (3 to 5 µm tip diameter) connected to a Picospritzer. Analyses of ΔF/F0 responses were performed in Python. The fluorescence (F) for each time point encompasses the averaged pixel intensities of the entire 512 x 512 image. Baseline fluorescence (F0) represents the averaged fluorescence values captured during the 1.5 s window before puffing. ΔF/F0 was calculated as (F-F0)/F0 for each time point to generate time series plots. Peak ΔF/F0 values were then extracted from the time series after puffing and plotted. To generate representative heatmaps, peak fluorescence for each pixel was averaged from a 3.75 s window following the puff. ΔF/F0 was then calculated for each pixel and plotted in the heatmap images. The resulting images were smoothened with a Gaussian blur (σ = 1).

### Stereotaxic Surgeries

Stereotaxic surgeries were performed following previously established methodology^14,16,20,23^. Mice were anesthetized with 5% isoflurane and mounted on a stereotaxic frame. 1.5 to 2% isoflurane was used to maintain stable anesthesia. Following exposure of the skull, a hand drill was used to create a small burr hole for syringe placement. For unilateral and bilateral injections in the dorsal striatum (coordinates relative to bregma: 1.0 mm anterior, ±2.0 mm lateral, 2.5 mm below the dura), adeno-associated viruses (AAVs) were delivered using a syringe coupled to a microinjector pump at a volume of 1 µL with a flow rate of 100 nL/min. Viruses were diluted to a working titer of 10^11^ copies/mL before injection. Mice used for fiber photometry were unilaterally equipped with fiberoptic cannulas (400 µm diameter) in the right dorsal striatum that were placed immediately following AAV delivery (coordinates relative to bregma: 1.0 mm anterior, 2.0 mm lateral, 2.3 mm below the dura) and fixed using quick adhesive dental cement. For unilateral and bilateral viral injections in M1 cortex (coordinates relative to bregma: 1.0 mm anterior, ±1.4 mm lateral, 1.25 mm below the dura) and the LC (coordinates relative to bregma: 5.40 mm posterior, ±1 mm lateral, 3.3 mm below the dura), manually pulled borosilicate glass pipettes (tip diameter 3 to 5 µm) were used for AAV delivery. A volume of 500 nL was delivered at a flow rate of 50 nL/min in M1. For bilateral AAV expression targeted to the LC, 1 µL of virus was injected at a flow rate of 100 nL/min on each side (coordinates relative to bregma: 5.40 mm posterior, ±1 mm lateral, 3.3 mm below the dura). For 6-hydroxydopamine hydrobromide (6- OHDA) injections in the right LC (coordinates relative to bregma: 5.40 mm posterior, 0.75 mm lateral, 3.3 mm below the dura), 1 µL of 6-OHDA (10 µg/µL in PBS containing 0.01% ascorbic acid) was delivered at a flow rate of 100 nL/min with a manually pulled borosilicate glass pipette (tip diameter 3 to 5 µm). Following injections, microinjectors were left in place for 10 minutes to allow the injected volume to settle. Mice were treated for postsurgical pain and were returned to their home cages for a minimum of 14 days before experiments.

### Adeno-associated viruses

AAVs were of the AAV9 serotype defined by the capsid, serotypes are included in figures independent of pseudotyping, and AAVs are available from Biohippo, BrainVTA, WZBioscience and Addgene. AAV9-hSyn-GRABDA2m (stocks at 1-2 x 10^12^ copies/mL, BioHippo, #BHV12400556-9) and AAV9-hSyn-GRAB_NE_2h (stocks at 1-2 x 10^12^ copies/mL, BioHippo, #BHV12400441-9) were purchased with permission from Y.L. AAV9-hSyn-nLightG (stocks at 1.4 x 10^12^ copies/mL, produced by the Viral Vector Facility of the University and ETH Zürich, AAV2/9.hSyn.nLightG) was provided by T.P. For photometry, AAV9-hSyn-GRAB_NE_2h was co-injected with AAV9-CAG-tdTomato (stocks at 1-2 x 10^13^ copies/mL, Addgene, #59462-AAV9).

### Slice imaging and analyses

Imaging in acute brain slices was performed as previously established^16,23^. Specifically, mice were deeply anesthetized with isoflurane and decapitated. The brain was dissected out and 250 μm thick brain slices were cut using a vibratome in ice-cold cutting solution containing (in mM) 75 NaCl, 2.5 KCl, 7.5 MgSO4, 75 Sucrose, 1 NaH2PO4, 12 Glucose, 26.2 NaHCO3, 1 Myo-inositol, 3 Na Pyruvic acid, and 1 Na Ascorbic acid. Sagittal slices were prepared for imaging in M1 cortex or dorsal striatum while coronal slices were cut for imaging in the LC. For experiments involving 6-OHDA ablation of the LC, sagittal slices from both hemispheres were cut. After cutting, slices were incubated for 30 minutes at 37 °C in a recovery solution composed of (in mM) 126 NaCl, 2.5 KCl, 2 CaCl2, 1.3 MgSO4, 1 NaH2PO4, 12 glucose, 26.2 NaHCO3, 1 Myo-inositol, 3 Na Pyruvic acid, and 1 Na Ascorbic acid. Slices were then incubated at minimum for 30 minutes at room temperature. Imaging was performed in a chamber continuously perfused with artificial cerebral spinal fluid (ACSF) containing (in mM) 126 NaCl, 2.5 KCl, 2 CaCl2, 1.3 MgSO4, 1 NaH2PO4, 12 glucose, and 26.2 NaHCO3 at 33 to 35 °C with a flow rate of 2 to 3 mL/min. All solutions were constantly bubbled with 95% O2 and 5% CO2, and data acquisition was completed within 5 hours of slicing. For Supplemental fig. 4, 1 µM of dihydro-β- erythroidine hydrobromide (DHβE, diluted from 100 mM stock) was included in the recording solution. Fluorescence imaging was performed with an Olympus BX51WI epifluorescence microscope. A 470 nm LED was used for excitation and the measured signals were digitized with a sCMOS camera (Hamamatsu Orca-Flash4.0). A 4x, 0.13 numerical aperture air objective was used for striatal imaging, while LC and M1 cortex imaging were performed with 10x, 0.30 numerical aperture or 40x, 0.80 numerical aperture water-immersion objectives, respectively. Electrical stimulation was applied using a pulled glass pipette filled with ACSF (3 to 5 µm tip diameter, 0.5 to 1 MΩ). Single stimuli and 20-stimulus 20 Hz trains were delivered at an intensity of 90 µA for 2 to 3 replicates with 2-minute interstimulus intervals using a linear stimulus isolator. A biphasic wave (0.25 ms in each phase) was applied for each stimulus. For Supplemental fig. 2a-d, fluorescence changes in responses to single and train stimuli were assessed before and after wash-on of tetrodotoxin (TTX, 1 µM) for 10 minutes. For Supplemental fig. 2e-h, fluorescence changes in responses to single and train stimuli were assessed before and after wash-on of a cocktail containing 6-Cyano-7-nitroquinoxaline-2,3- dione (CNQX, 10 µm), D-AP-5 (20 µm), SR-95531hydrobromide (gabazine, 10 µM), CGP- 55845 hydrochloride (300 nm), atropine (30 nm), and DHβE (1 µm) for 20 minutes. For Supplemental fig. 2i-l, fluorescence changes in responses to single and train stimuli were assessed before and after incubation in ACSF for 20 minutes. Fluorescence images were acquired at 512 x 512 pixels/frame and 50 frames/s with an exposure time of 20 ms. Striatal slice imaging using DHβE was performed at 20 frames/s with an exposure time of 50 ms per frame. Image analyses were performed in Python. For quantification of transients in the dorsal striatum and LC (Figs. 1 and 2, and Supplemental figs. 4, 6, and 8), circular regions of interest (ROIs) with 30- to 50-pixel radii were manually selected in each image time series to calculate time series of ΔF/F0. The position of each ROI was centered around the visually identified peak signal. The radii of manually selected ROIs were constant for each brain area at 50 pixels for LC and 30 pixels for dorsal striatum. For recordings in M1 cortex shown in Fig. 1 and Supplemental. fig. 2, ROIs were generated from the top 10% of responding pixels in the first 20-stimuli replicate. The resulting mask was then used to calculate ΔF/F0 in the remaining replicates. M1 cortex recordings in Fig. 3 were analyzed using manually selected circular ROIs with a radius of 100 pixels. For recordings in M1 cortex, LC, and dorsal striatum with nLightG or in the presence of DHβE, bleach correction was performed on raw signals with Python’s SciPy’s curve fitting module using an exponential decay function. To determine baseline fluorescence (F0), 500 ms windows before the onset of electrical stimulation were averaged within an ROI. The fluorescence (F) for each time point represents the averaged pixel intensities within each ROI.

ΔF/F0 was then calculated as (F-F0)/F0 for each time point and averaged to generate the time series plots. Peak ΔF/F0 values were then extracted from the time series and plotted. This process was also used to create example heat plot images for the time window 280-500 ms after stimulation. The resulting images were smoothened with a Gaussian blur (σ = 1) to generate the example heat plot images for display in the figures. Recordings from control and experimental conditions were processed identically.

### Fiber photometry

Fiber photometric recordings in freely moving mice were performed as previously established^16,23^. At 21 days after surgical implantation, the fiberoptic cannula was connected to an optic patch cord, and the mice were allowed to freely roam for 1-hour epochs in a circular arena (43.1 cm diameter, 35.6 cm high) illuminated by infrared light (850 nm, 30 μW/cm^2^ as measured from the arena surface). Silicon photodiodes were used to convert fluorescence to electrical signals, which were then amplified by photodiode amplifiers and collected by a multifunction I/O card at 10,000 Hz. 470 nm and 565 nm LEDs (Thorlabs) were used to excite the channels and applied at 157 μW and 17 μW, respectively, and as established before^16^.

Output power was chosen to hold the raw fluorescence signals for green and red channels in a similar range, and excitation for each LED was applied in an alternating pattern at 25 Hz. During the 40 ms active period, each excitation channel was on for 10 ms and the average output of each channel was assessed and set according to previously established protocols^23^. For analyses, the raw photometry signal F was first processed with a low-pass filter at 0.01 Hz to estimate F0, and ΔF/F0 was calculated as (F-F0)/F0 and analyzed from the first 30 minutes of the 1-hour data acquisition. To quantify the fluorescence fluctuations, a 640 ms sliding window was applied to ΔF/F0 transients to calculate an average of the standard deviation. Brief illuminations (200 ms light pulses, 565 nm LED light source, 50 μW/cm^2^ measured at the bottom of the arena) were applied at random time intervals ranging from 60 to 210 s during the 1 h epoch. At the end of the experiment, animals were deeply anesthetized with 5% isoflurane and transcardially perfused with 30 mL PBS followed by 40 mL 4% PFA in PBS. Brains were then extracted and post-fixed in 4% PFA in PBS for 24 hours following perfusion. Brain slices were cut at 50 µm in ice-cold PBS on a vibratome. Sections were mounted on glass slides with a mounting medium containing DAPI. Images of the brain sections were acquired using a slide scanning microscope with a 4x air objective. Cannula positions were determined with respect to a mouse brain atlas^38^ and are displayed in Supplemental fig. 5 on schematics drawn from corresponding brain sections.

## Statistics

Data are shown as mean ± SEM with * p < 0.05, ** p < 0.01, *** p < 0.001. Data were plotted using Python (Python Software Foundation, 3.11.4), MATLAB (MathWorks R2022b) and/or Prism (GraphPad 10.2.3), and individual data points are included in each figure whenever possible. The number of observations in each experiment was based on previous publications^14,16,23,39^ and on trial experiments; no statistical methods were used to predetermine sample sizes. For each experiment, the definitions of n are specified in the figure legends, and sample sizes for each plot are included in each figure legend. For genotype comparisons, the experimenter was blind to genotype while acquiring and analyzing data. Data sets were tested for normality with Anderson-Darling, D’Agostino-Pearson omnibus, Shapiro-Wilk, and Kolmogorov-Smirnov tests. Data sets were tested for variance with the F test for two-group comparisons and the Brown-Forsythe and Bartlett’s tests for three-group comparisons. Data sets that met criteria for normalcy and equal variance across all tests were analyzed using parametric statistical tests. Non-parametric tests were used otherwise. Specifically, two-tailed Mann-Whitney rank-sum tests were used for Figs. 1c, 2e, 2h, 3c, 3g, 3k, and 3o (single stimulus), and Supplemental figs. 3c, 4d, and 7c. Two-tailed Wilcoxon sign-rank tests were used for Figs. 1g and 1j, and Supplemental figs. 2d, 2h, 8d, and 8g. Two-tailed paired t-tests were used for Supplemental fig. 2l. Two-tailed unpaired t-tests were used for Figs. 2k and 3o (20 stimuli), and Supplemental fig. 6f. One-way ANOVA and Dunnet’s multiple comparisons post-hoc tests were used for Supplemental fig. 2m. Kruskal-Wallis and Dunn’s multiple comparisons tests were used for Supplemental fig. 2n. The specific statistical tests used are also described in each figure legend.

---

**Supplemental figure 1.**
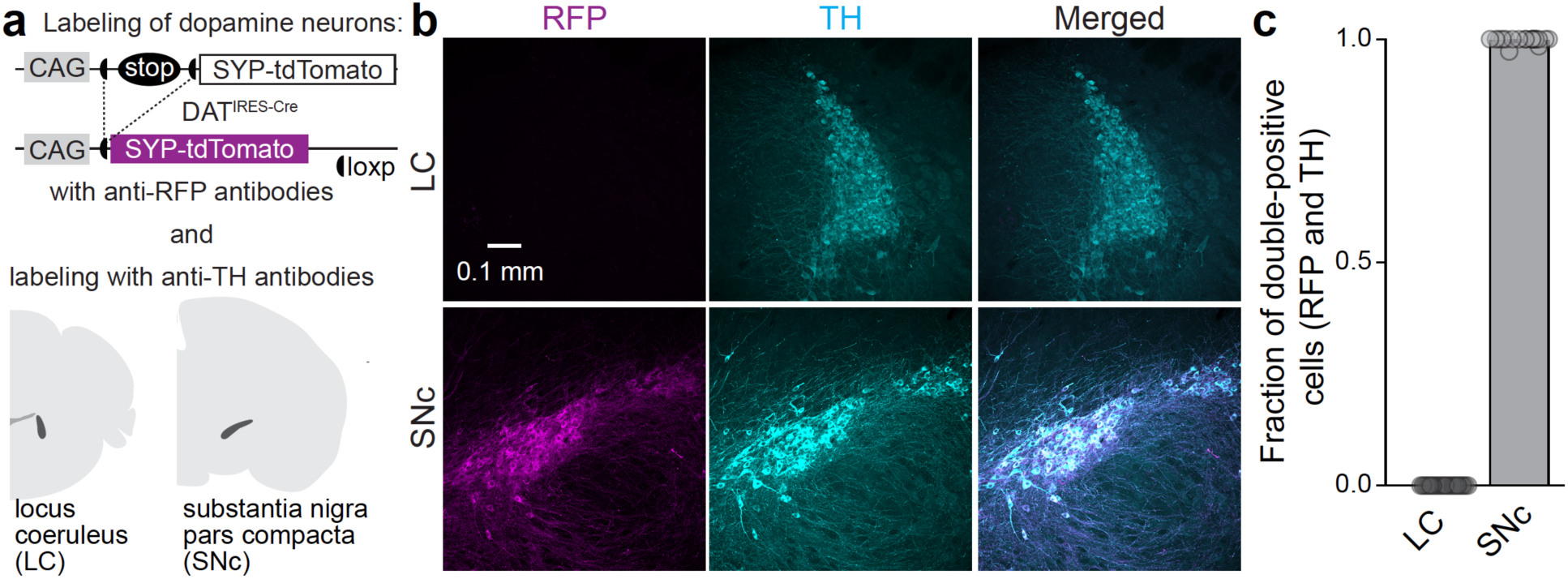
Assessment of Cre-dependent expression of synaptophysin- tdTomato in DAT^IRES-Cre^ mice. **a.** Strategy for labeling dopamine and norepinephrine neurons similar to Fig. 1a-c but with staining for tyrosine hydroxylase (TH) followed by analyses of immunofluorescence in locus coeruleus (LC) and substantia nigra pars compacta (SNc). **b, c.** Representative confocal images of LC and SNc (b) and quantification of the fraction of cells double positive for RFP and TH (c); LC 20 slices from 4 mice, SNc 20/4. Data are mean ± SEM. On average, 96 ± 6 cells in LC and 88 ± 6 cells in SNc were positive for TH per image. The observation that there are no double-positive cells in LC indicates that the DAT^IRES-Cre^ line does not express Cre in LC neurons. Hence, the strategy of NET labeling combined with RFP-labeling of SYP-tdTomato in DAT^IRES-Cre^ mice can be used to distinguish norepinephrine from dopamine axons. Both norepinephrine and dopamine neurons express TH.

**Supplemental figure 2.**
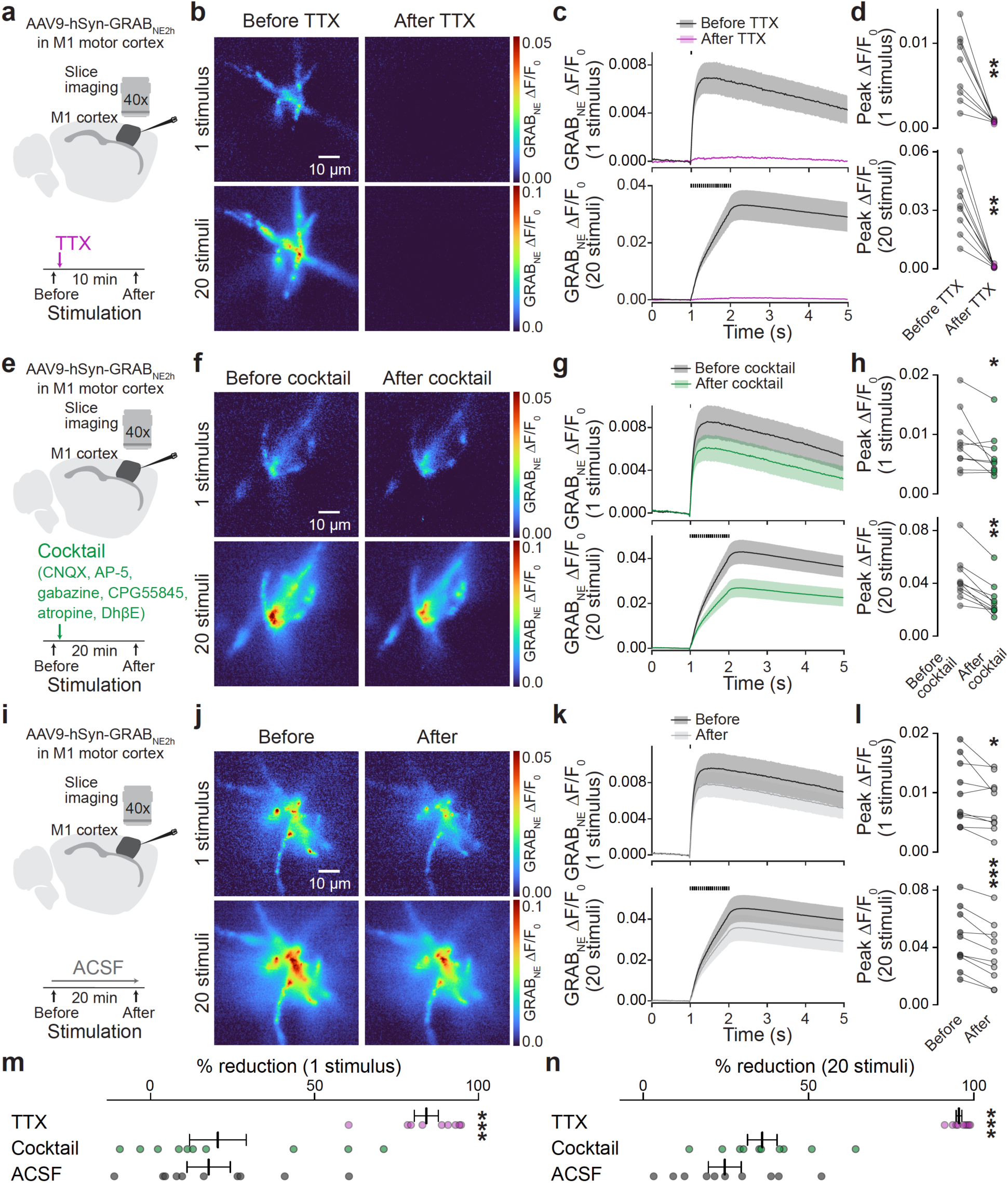
Pharmacological inhibition of GRAB_NE_ transients in brain slices. **a.** Schematic of imaging in cortex before and after 10-minute bath application of the sodium channel blocker tetrodotoxin (TTX) to inhibit action potential firing. **b-d.** Representative images of peak GRAB_NE_ fluorescence in response to 1 stimulus or 20 stimuli at 20 Hz before and after TTX wash-on (b) and quantification of GRAB_NE_ ΔF/F0 time series (c) and peak ΔF/F0 (d); 9 slices from 3 mice. **e-h**. Same as in a-d, but with bath application of a drug cocktail containing CNQX, AP-5, gabazine, CGP-55845, atropine, and DHβE for 20 min; 10/3. **i-l**. Same as in a-d, but with incubation in ACSF for 20 min; 10/3. **m, n.** Post-hoc assessment of percent reduction in response to 1 stimulus (m) or 20 stimuli (n) comparing incubation in TTX (a-d, 10 min), the drug cocktail (e-h, 20 min), or ACSF (i-l, 20 min); TTX 9/3, cocktail 10/3, and ACSF 10/3. Overall, rundown during drug cocktail application and incubation in ACSF was similar while TTX strongly inhibited the fluorescence changes. Data are mean ± SEM; *** p < 0.001, ** p < 0.01, * p < 0.05, as assessed by: two-tailed Wilcoxon signed-rank tests in d and h; two-tailed paired t-tests in l; one-way ANOVA and Dunnet’s multiple comparisons post-hoc tests in m (compared to ACSF); Kruskal-Wallis and Dunn’s multiple comparisons tests in n (compared to ACSF).

**Supplemental figure 3.**
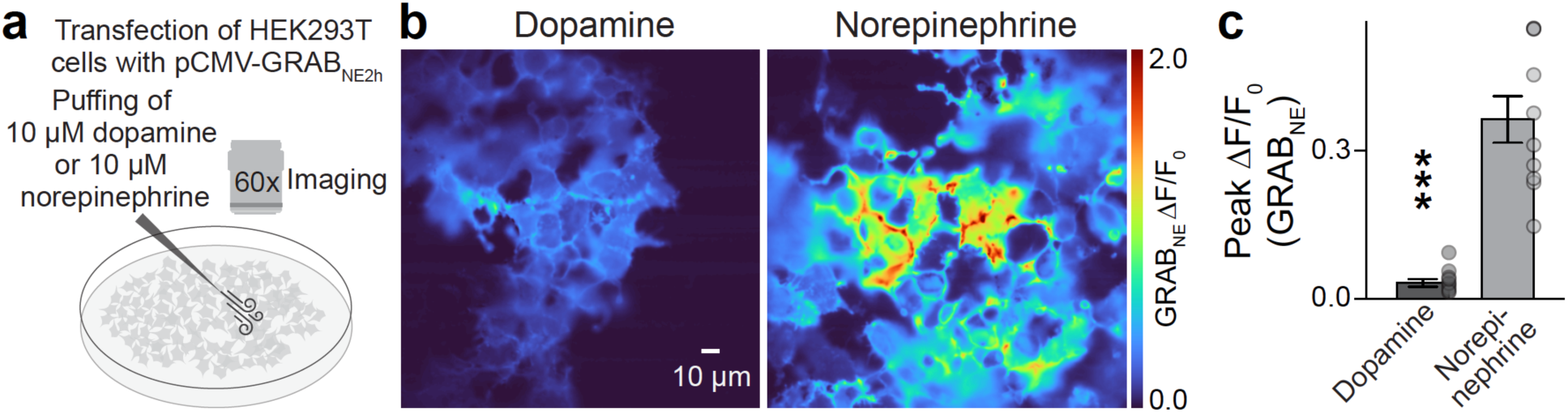
Assessment of GRAB_NE_ fluorescence in HEK293T cells. **a.** Schematic of the experiment in which 10 µM of dopamine or norepinephrine were puffed onto HEK293T cells transfected with pCMV-GRAB_NE_2h. **b, c.** Representative peak GRAB_NE_ fluorescence (b) and peak ΔF/F0 (c); 10 coverslips from 3 transfections each. Data are mean ± SEM; *** p < 0.001, as assessed by two-tailed Mann-Whitney rank-sum test in c.

**Supplemental figure 4.**
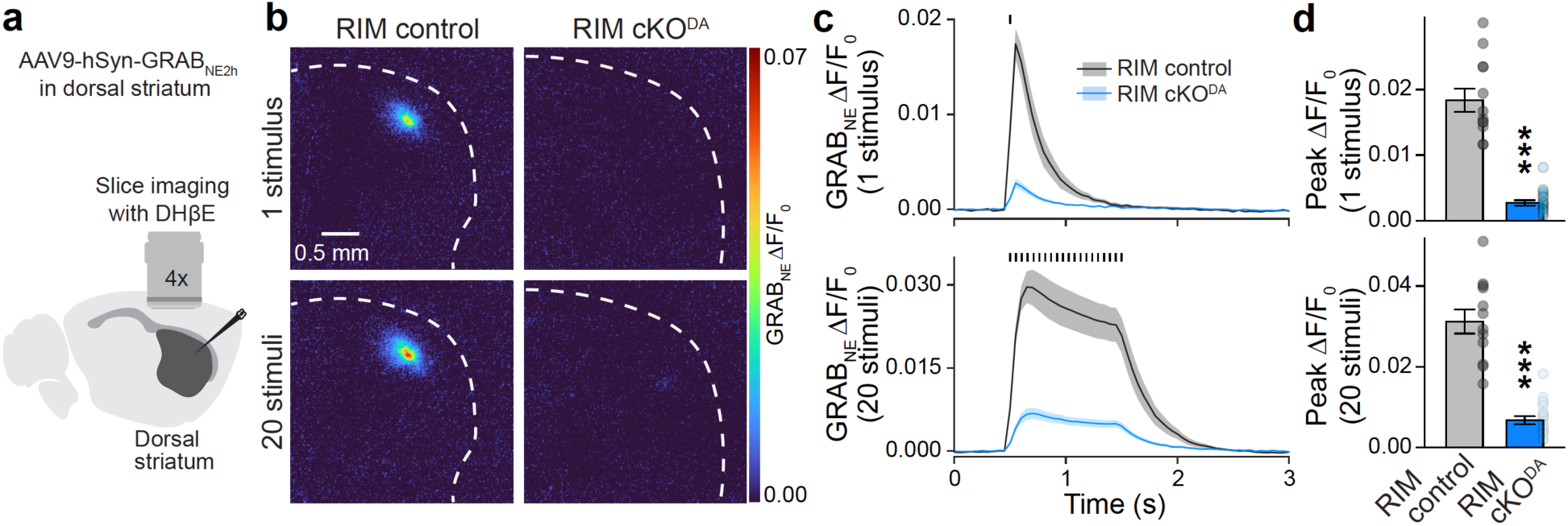
GRAB_NE_ recordings in striatum in the presence of DhβE. **a.** Schematic of imaging experiments as in Fig. 2c-e, but in the presence of dihydro-β-erythroidine hydrobromide (DHβE, 1 µM) to block β2-containing nicotinic acetylcholine receptors. **b-d.** Representative images of peak GRAB_NE_ fluorescence in response to 1 stimulus or 20 stimuli at 20 Hz (b) and quantification of GRAB_NE_ ΔF/F0 time series (c) and peak ΔF/F0 (d); RIM control 12 slices from 3 mice, RIM cKO^DA^ 17/4. Data are mean ± SEM; *** p < 0.001, as assessed by two-tailed Mann-Whitney rank-sum tests in d.

**Supplemental figure 5.**
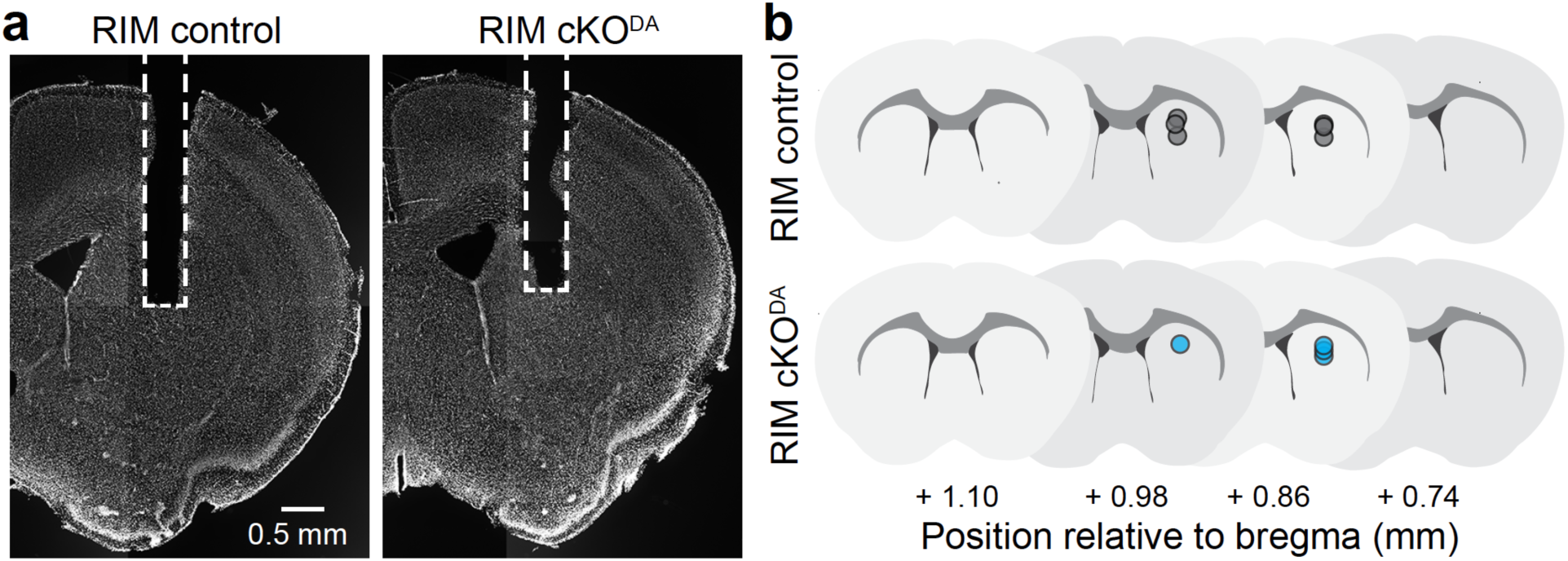
Fiberoptic cannula positions. **a.** Representative images of coronal sections of mice with fiberoptic cannulas. Coronal sections were imaged with a slide scanner in DAPI-containing mounting medium. **b.** Illustration of fiberoptic cannula tip positions of the mice analyzed in Fig. 2k determined relative to a mouse brain atlas^38^ and mapped onto schematics drawn from brain sections.

**Supplemental figure 6.**
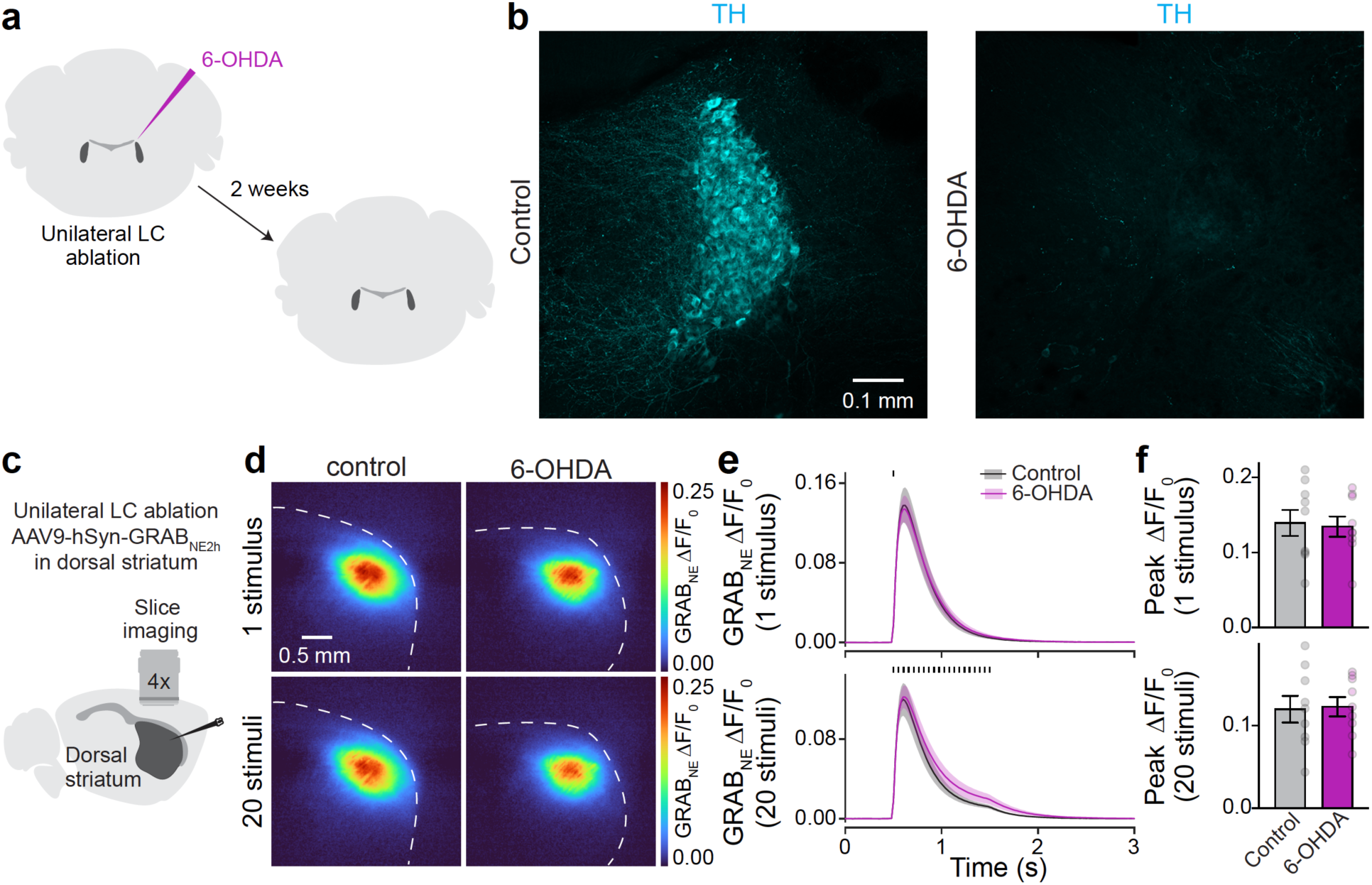
Striatal GRAB_NE_ responses after 6-OHDA lesion of LC. **a.** Schematic of unilateral 6-OHDA ablation of LC followed by assessment with confocal microscopy. **b.** Confocal images of LC after staining with TH antibodies; a representative image of 4 slices from 4 mice is shown. **c.** Schematic of bilateral slice imaging in dorsal striatum expressing GRAB_NE_ with unilateral 6- OHDA ablation of the right LC. **d-f.** Representative images of peak GRAB_NE_ fluorescence in response to 1 stimulus or 20 stimuli at 20 Hz in ipsilateral (6-OHDA) and contralateral (control) hemispheres (d) and quantification of GRAB_NE_ ΔF/F0 time series (e) and peak ΔF/F0 (f); control 9 slices from 3 mice, 6-OHDA 9/3.

**Supplemental figure 7.**
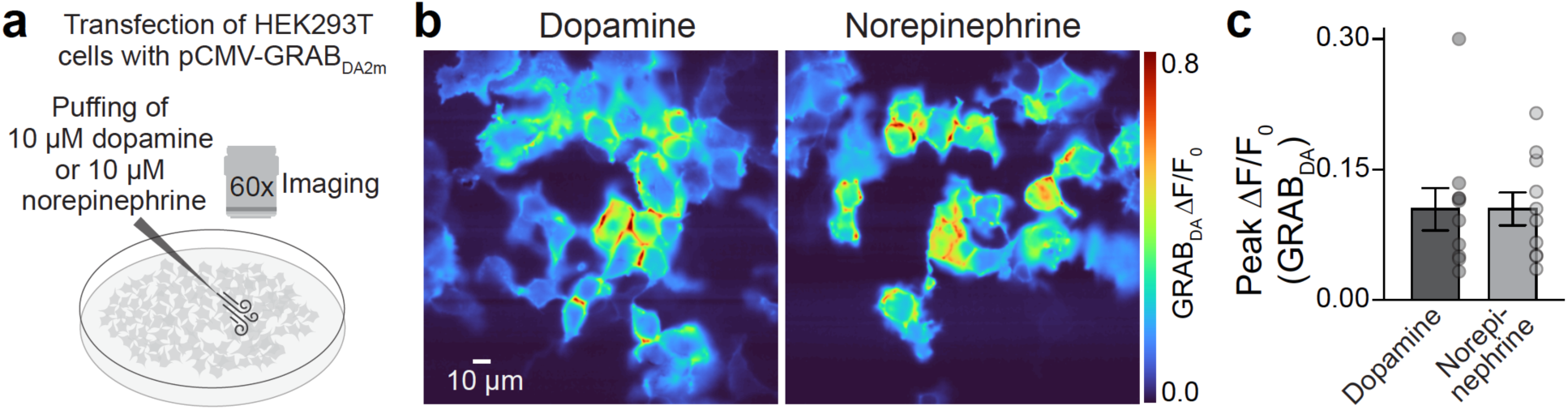
Assessment of GRABDA fluorescence in HEK293T cells. **a.** Schematic of the experiment in which 10 µM of dopamine or norepinephrine were puffed onto HEK293T cells transfected with pCMV-GRABDA2m. **b, c.** Representative peak GRABDA fluorescence (b) and peak ΔF/F0 (c); 10 coverslips from 3 transfections each. Data are mean ± SEM.

**Supplemental figure 8.**
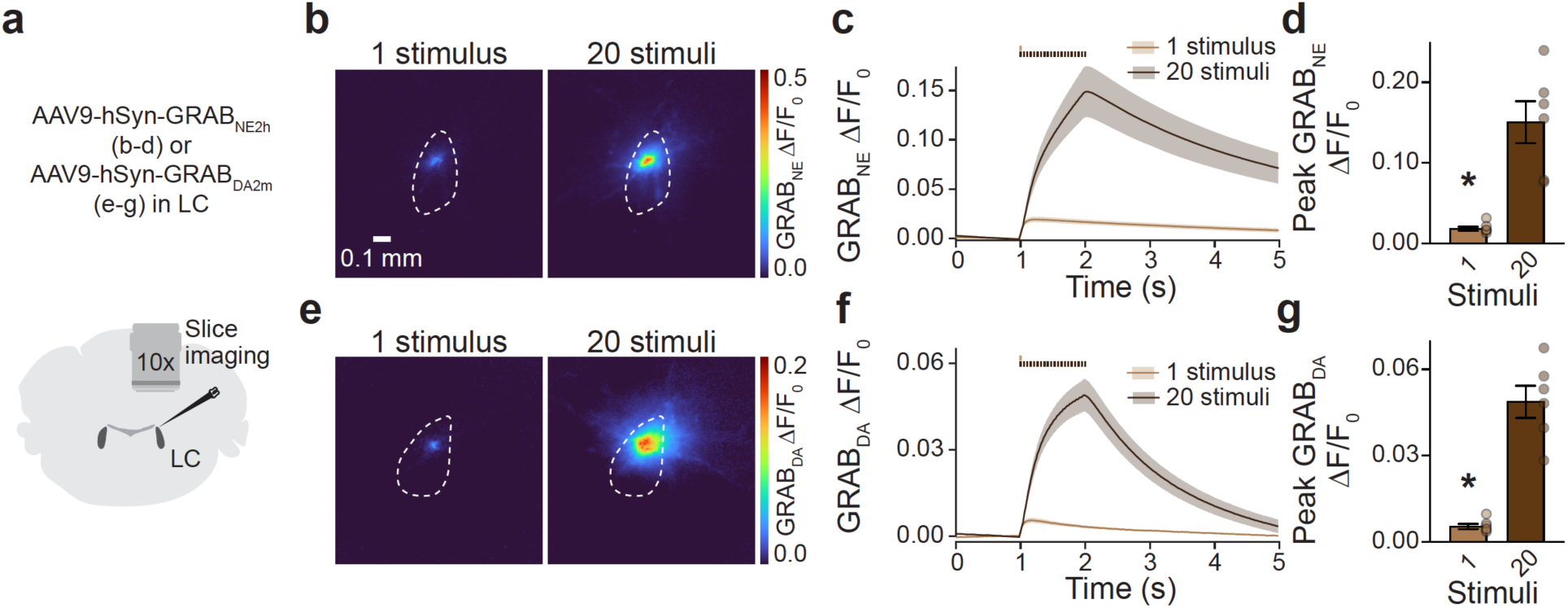
Norepinephrine and dopamine sensors in LC. **a.** Schematic of slice imaging in LC. **b-d.** Representative GRAB_NE_ peak fluorescence in response to 1 stimulus or 20 stimuli at 20 Hz in LC (b) and quantification of ΔF/F0 time series (c) and peak ΔF/F0 (d); 6 hemispheres from 3 slices from 3 mice. **e-g.** As in b-d, but with GRABDA; 6/3/3. Data are mean ± SEM; * < 0.05 as assessed by two-tailed Wilcoxon signed-rank tests in d and g.

## Supplemental text. Studies with GPCR-based sensors in brain areas with dual dopamine and norepinephrine innervation

In the present study, we tested whether GPCR-based dopamine and norepinephrine sensors are specific for their corresponding transmitters in the brain. These sensors have been widely used in brain areas that are innervated by both norepinephrine and dopamine neurons and have been expressed in mice and rats.

Previous studies have used norepinephrine or dopamine sensors in the cortex^3,4,40–50^ and in the cerebellum^51^, brain areas in which norepinephrine innervation is typically higher in density compared to dopamine innervation^7,10^ (Fig. 1a-c). Areas with dopamine and norepinephrine innervation densities that are likely in a similar range^52–56^ have also been studied with these sensors, for example the hippocampus^5,57–60^, the basolateral amygdala^3,49^, the lateral hypothalamus^4,60–62^, and the basal forebrain and thalamus^63^. Finally, dually innervated brain areas in which dopamine dominates over norepinephrine^55,64,65^ have been characterized with the sensors as well, for example the ventral striatum^1–3,16,49,50,62,66–72^ and the ventral tegmental area^49^.

Overall, the sensor signals in these brain regions might be due to either dopamine or norepinephrine release. Without cell type- or projection-specific manipulations to remove either dopamine or norepinephrine innervation and/or secretion, it is difficult to conclude which transmitter is detected in dually innervated brain areas when GPCR-based sensors for dopamine or norepinephrine are used.

